# Evaluating Few-Shot Meta-Learning using STUNT for Microbiome-Based Disease Classification

**DOI:** 10.64898/2026.03.01.708821

**Authors:** Chengyao Peng, Thomas Abeel

**Affiliations:** Delft Bioinformatics Lab, Delft University of Technology, Delft, The Netherlands; Infectious Disease and Microbiome Program, Broad Institute of MIT and Harvard, Cambridge, USA

## Abstract

The human gut microbiome is increasingly explored as a diagnostic indicator for disease, yet machine learning models trained on metagenomic data are often constrained by limited sample sizes and poor cross-cohort generalizability. Meta-learning, a machine learning paradigm that optimizes models for rapid adaptation to new tasks with limited examples, offers a promising strategy to address this by leveraging the potential shared microbial structure across publicly available metagenomic datasets. Here, we evaluated STUNT, a framework combining self-supervised pretraining with metric-based meta-learning (Prototypical Networks), for few-shot microbiome-based disease classification. Using over 5,000 species-level gut metagenomic profiles from 57 cohorts in GMrepo v2, we meta-trained STUNT on 52 cohorts and evaluated the pretrained embedding on five held-out disease cohorts covering rheumatoid arthritis (RA), gestational diabetes mellitus during pregnancy (GDM), non-alcoholic fatty liver disease (NAFLD), diabetes mellitus, type 1 (T1D), and inflammatory bowel disease (IBD). We compared Prototypical Networks, Logistic Regression, and Random Forest with and without STUNT-derived embeddings across shot sizes of 1 to 10 samples per class. We found that STUNT-derived embeddings provided a modest benefit only under extreme data scarcity (one labeled sample per class) and this advantage rapidly diminished and reversed with additional samples, indicating that the meta-learned representations impose an information bottleneck limiting access to task-specific signals. Classification performance varied substantially across cohorts, consistent with PERMANOVA-estimated microbiome–disease separability. These results highlight the need for representation learning approaches that preserve disease- and cohort-specific variation and suggest that intrinsic biological signal strength is the primary determinant of classification success.

## 1 Introduction

The gut microbiome is closely involved in host metabolism and health. While shaped by various host and environmental factors, including host genetics, diet, and lifestyle, the human gut microbiome in turn regulates human physiological and immune processes through microbial metabolism and signaling [1]. Changes in the human gut microbiome can therefore both reflect and drive disease processes. As a result, there is growing interest in understanding the role of the human gut microbiome and its links to a wide range of human diseases, including intestinal, metabolic, immunological, and neurological disorders [2, 3].

With the increasing availability of metagenomic sequencing data, machine learning methods, especially supervised algorithms, have been used to associate gut microbiome compositions with different human conditions [4]. Although these models have shown promising predictive performance within individual datasets, their generalizability between cohorts is frequently constrained [5]. This limitation stems primarily from the intrinsic characteristics of microbiome data: compositionality, high dimensionality, and sparsity, where the number of microbial taxa far exceeds the number of samples [6]. Moreover, substantial inter-individual variability in microbiome composition, combined with its susceptibility to biological and technical confounders, further increase the risk of overfitting and poor generalizability of machine learning models across different cohorts [7, 8].

Recent advances in self-supervised representation learning and transfer learning have transformed multiple areas of computational biology. For example, protein language models trained with self-supervised objectives on large-scale protein sequence databases can learn representations that encode protein structural and functional information and support related downstream prediction tasks [9]. In transcriptomics, large-scale representation models demonstrate that self-supervised pretraining can learn transferable gene-expression representations for diverse downstream genomic tasks, such as cell-type classification and perturbation prediction [10]. These approaches aim to learn general feature mappings from unlabeled data, in other words, without requiring additional task-specific annotations. The resulting pretrained representations can then be reused through transfer learning, in which the pretrained model is adapted or fine-tuned for a specific prediction task using relatively limited labeled data. Such a strategy is also increasingly being explored in tasks linking the gut microbiome with human diseases to overcome limitations imposed by the intrinsic characteristics of microbiome data [11–13]. Beyond self-supervised pretraining, transfer learning alone has also been used to address cross-cohort variability in microbiome-based disease classification. For example, Wang et al. [14] applied transfer learning to improve cross-regional microbiome-based disease diagnosis by adapting supervised models trained in one geographical location to another. Collectively, these studies demonstrate that pretrained representations can potentially mitigate data scarcity and cross-study variability. However, these approaches typically rely on a single pretraining objective and need task-specific fine-tuning for downstream tasks.

Given the recent meta-analyses suggesting the existence of generalizable patterns in human gut microbiome data across populations [15] and diverse conditions [16–18], methods and strategies that are able to explicitly leverage this shared microbial structure or latent information across cohorts may further improve model adaptation efficiency. One such advanced approach is meta-learning, which builds upon principles in transfer learning but extends them in key ways. Compared to transfer learning, meta-learning explicitly learns model parameters by leveraging the shared structure across multiple related tasks, while simultaneously optimizing the model for rapid adaptation to new tasks [19]. In domains such as computer vision and natural language processing, meta-learning has shown superior performance over standard supervised and transfer learning approaches in scenarios involving multiple related tasks, each characterized by limited data [20, 21]. This formulation is well suited to microbiome-based disease classification, where multiple related classification tasks naturally arise across cohorts, populations, and disease contexts, while labeled data within each classification task is often limited, though the extent to which shared structure is learnable across heterogeneous cohorts remains unclear.

Metric-based meta-learning approaches, such as Prototypical Networks, enable efficient meta-learning by learning a shared prototypical embedding space. In this space, samples can be classified according to their distances to class prototypes, constructed from only a few support examples [22]. STUNT (Few-shot Tabular Learning with Self-generated Tasks from Unlabeled Tables), combines the advantages offered by self-supervised representation learning with metric-based meta-learning [23]. Specifically, STUNT uses self-supervised pretraining to learn feature representations from unlabeled data, and then apply a Prototypical Network to leverage these representations for new tasks. Currently in microbiome research, although raw sequencing data are often deposited in public repositories, processed data, phenotypic metadata, and computational workflows are frequently incomplete or unavailable [24]. Therefore, STUNT provides a suitable framework for learning task-adaptable representations in microbiome-based classification settings, where high-quality labeled data are scarce.

In this study, we systematically assess whether STUNT can improve microbiome-based disease classification with limited labeled data availability. Using GMrepo v2 [25], a curated repository of human gut metagenomic datasets, we construct a meta-learning framework that includes 52 diverse projects for meta-training and five held-out disease cohorts for evaluation, covering inflammatory bowel disease, type 1 diabetes, rheumatoid arthritis, non-alcoholic fatty liver disease, and gestational diabetes mellitus. We evaluated the meta-learned embedding by comparing the performance of Prototypical Networks, Logistic Regression and Random Forest trained with and without leveraging the STUNT-derived embeddings, under both few-shot and full-data settings. Our results show that the pretrained embedding provides a modest advantage only under extreme data scarcity (one labeled sample per class), but this benefit diminishes and reverses as more labeled samples become available. Furthermore, classification performance varies substantially across disease cohorts, consistent with PERMANOVA-estimated separability of microbial profiles between case and control groups. Our work outlines priorities for future research on pretrained models for microbiome-based disease classification, helps clarify the conditions under which self-supervised representation learning is likely to succeed or fail, and underscores the importance of examining intrinsic biological signal strength when evaluating methods developed in this domain.

## 2 Methods

We evaluated STUNT, a self-supervised meta-learning framework, for its ability to learn generalizable patterns in human gut microbiome data and improve microbiome-based disease prediction under limited labeled data. An overview of the workflow, including data pre-processing, meta-training, meta-testing, and baseline evaluation, is summarized in Figure 1, with each component described in detail below.

**Figure 1:**
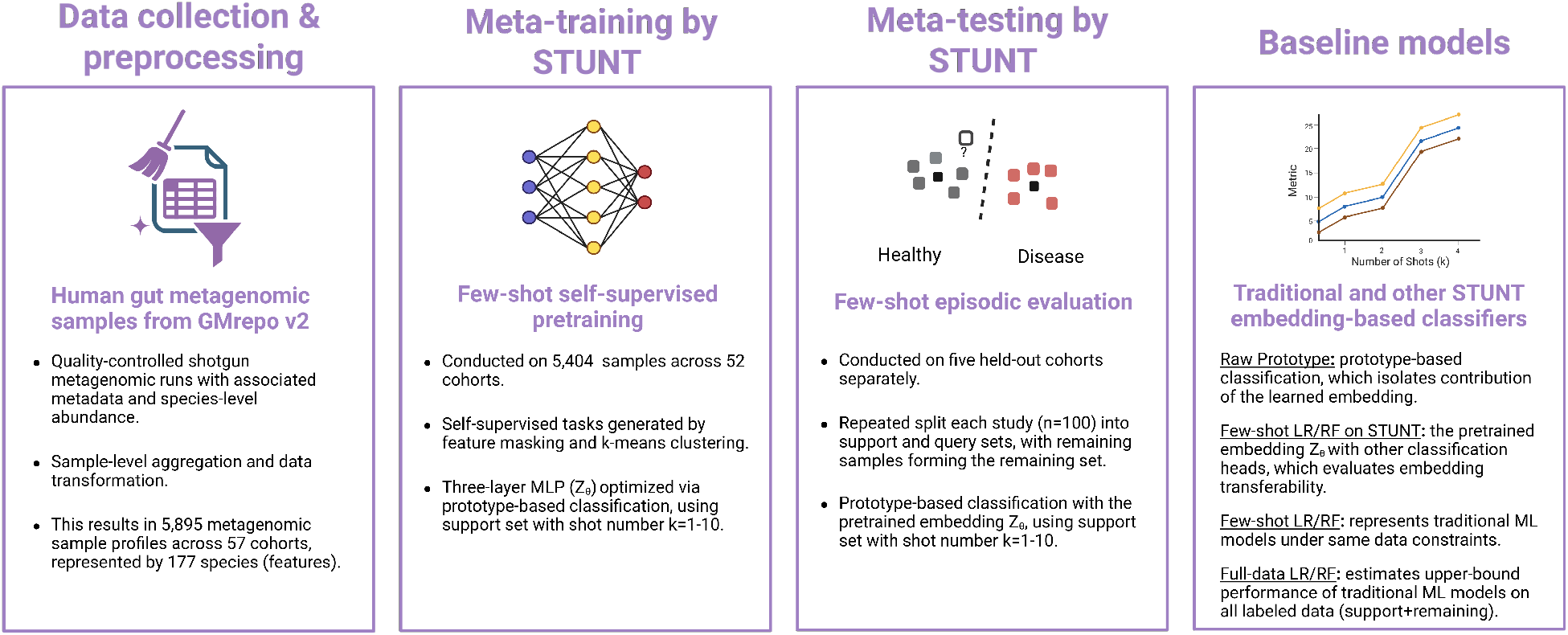
Overview of the STUNT framework and evaluation pipeline for few-shot microbiome-based disease classification. To prepare the metagenomic data used to train and evaluate STUNT and baseline methods, we collected and preprocessed 5,895 human gut metagenomic profiles across 57 cohorts from GMrepo v2, represented by 177 microbial species abundances. In meta-training, STUNT conducts few-shot self-supervised pretraining on 5,404 samples from 52 cohorts using feature masking and k-means–based task generation. During this phase, a three-layer MLP encoder is optimized by performing self-generated tasks through prototype-based classification across varying shot numbers (*K*=1–10). In later meta-testing phase, the pretrained STUNT embedding is evaluated via few-shot episodic classification on five held-out cohorts using repeated support–query splits (n=100 times) and prototype-based classification. The performance of STUNT is compared against several baseline models, which were trained only on the held-out studies and evaluated using the same repeated episodic splits as STUNT. These models include a Prototypical Network on raw features (Raw Prototype), Few-shot Logistic Regression/Random Forest on STUNT-derived embeddings (Few-shot LR/RF on STUNT), traditional LR/RF trained on raw features, instead of STUNT-derived embeddings (Few-shot LR/RF), and traditional LR/RF trained on all available labeled data using raw features, excluding the query set (Full-data LR/RF).

### 2.1 Parsing shotgun metagenomic samples from GMrepo v2

We parsed the species-level abundance of metagenomic samples provided by GMrepo v2 [25], one of the largest databases of curated and annotated human gut metagenomes. The database extensively aggregated human gut microbiome projects from NCBI BioProject (https://www.ncbi.nlm.nih.gov/bioproject/) and PubMed (https://pubmed.ncbi.nlm.nih.gov/), with raw reads downloaded from NCBI SRA and EBI ENA repositories. The associated metadata, including sequencing details, phenotypes, age, sex, BMI, and antibiotic use, were also collected if available and manually curated. The raw reads were quality-controlled and taxonomically profiled using a unified pipeline to facilitate cross-project analysis. We downloaded the taxonomic abundance data, curated metadata and project information provided at https://evolgeniusteam.github.io/gmrepodocumentation/usage/downloaddatafromgmrepo/, as accessed on 15 January 2025.

In total, GMrepo v2 contained 52,633 processed runs, including 16,156 shotgun metagenomic runs and 36,477 amplicon sequencing runs. To avoid technical artifacts from the two different sequencing techniques, we restricted the analysis to shotgun metagenomic runs since whole-genome shotgun metagenomic sequencing normally provides more resolution and has increasingly become the norm of human microbiome research. We further retained the runs that passed the quality control (13,643), have associated metadata (12,109) and have the species-level abundance data (11,569). These runs corresponded to 5,895 unique samples. When multiple runs corresponded to the same biological sample, their abundance profiles were summed and renormalized to obtain a single sample-level relative abundance profile. After pre-processing, the dataset comprised 5,895 samples spanning 57 cohorts.

### 2.2 Study selection for meta-training and testing

Since training and evaluating STUNT only requires labeled data during meta-testing, we first decided on projects for meta-testing and used the remaining projects for meta-training. Specifically, to emulate real-world cross-cohort prediction scenarios, we treated the disease classification task within each cohort as an independent meta-testing task.

To pick the projects used for meta-testing, we first excluded one project lacking disease phenotype annotation. We then restricted candidate projects to those containing between 20 to 200 samples, a range that is typical for human microbiome datasets. To maintain consistent binary classification settings, we excluded cohorts associated with more than two phenotypes and those with fewer than 15 samples in either class. After applying these criteria, five projects were retained as held-out cohorts for meta-testing. Samples from the remaining 52 projects were used for meta-training.

### 2.3 STUNT few-shot meta-training setup

To remove noise in the meta-training data, we filtered out species which are absent in more than 95% of the meta-training samples, leaving 177 species-level features for downstream analysis. We then applied a center-log-ratio (CLR) transformation to account for the compositionality, followed by z-score standardization to ensure that all taxa contributed comparably in the later distance-based learning.

During meta-training, STUNT generated self-supervised classification tasks from the unlabeled feature table for model training and validation. These tasks are generated on-the-fly with over 10,000 training steps and a task batch size of 4, resulting in 40,000 total tasks. Models were trained with shot numbers ranging from *K* = 1 to *K* = 10, where *K* denotes the number of support samples per class. All other hyperparameters were kept at their default settings.

For each generated task, a random subset of features was selected using a masking ratio of 20-50%. Based on these selected features, STUNT applied k-means clustering (*C* = 10) to all meta-training samples to generate pseudo-labels. From the pseudo-label data, a balanced and disjoint support set *S* and query set *Q* were sampled. The support set *S* contained 10 pseudo-classes with *K* samples per class (10-way *K*-shot), and the query set *Q* contained 10 pseudo-classes with 15 samples per class.

STUNT optimizes a task-adaptable embedding function through a prototype-based meta-learning objective, in which samples are mapped into a shared embedding space and classified according to their distance to class prototypes. The embedding function is parameterized as a three-layer multilayer perceptron (MLP) and optimized across the generated tasks. Class prototypes were computed as the mean embeddings of support samples within each pseudo-class. Let *f_θ_*(*·*) denote the embedding function parameterized by meta-parameters *θ*. For each pseudo-class *c*, the prototype **p***_c_* is defined as:

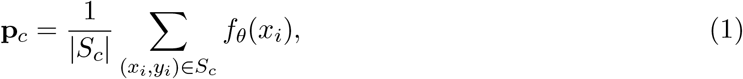

where *S_c_*denotes the subset of support samples belonging to class *c*. Query samples were then classified based on their Euclidean distance to the computed class prototypes and the probability of assigning a query sample *x* to class *c* is given by:

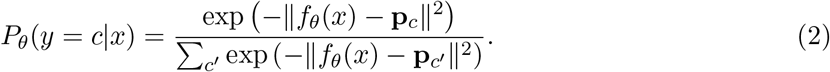

The embedding parameters *θ* were optimized during meta-training by minimizing the cross-entropy loss over query samples:

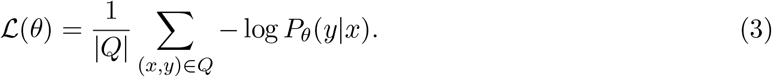

Gradients are back-propagated through the prototype computation and embedding function to update the meta-parameters *θ* after each task batch. Iterating over many self-generated tasks encourages the embedding function to learn representations that generalize to unseen classification problems. For each shot value *K* from 1 to 10, the best checkpoint was selected based on the highest average validation accuracy across the query sets.

### 2.4 STUNT few-shot meta-testing setup

For meta-testing, the five held-out cohorts were preprocessed using the same pipeline as in meta-training. Specifically, we retained the same 177 species features and performed CLR transformation, followed by z-score standardization. Prior to meta-testing, we used permutational multivariate analysis of variance (PERMANOVA) [26] to quantify the proportion of variance in microbiome composition explained by disease status in each cohort. This step provides a cohort-wise estimate of disease-associated signal strength, allowing us to contextualize the few-shot classification performance across studies with varying degrees of microbiome–disease separability.

To evaluate the benefit of using the meta-learned embedding in a few-shot setting, we adopted an episodic procedure similar to meta-training, but without updating the meta-parameters, *θ*. For each evaluation episode for each study, samples were randomly split into three disjoint sets: a support set *S*, a query set *Q* and a remaining set *R*. The support set was constructed by randomly selecting exactly *K* samples from each class. From the remaining samples, 10 were randomly selected to form the query set *Q*, and all other samples formed the remaining set *R*.

During evaluation, similar to meta-training, the pretrained embedding function *f_θ_* generated embeddings for the support samples in *S*, and class prototypes were computed as the mean embeddings for each class (Eq. 1). Each query sample in *Q* was then embedded and classified according to its Euclidean distance to the class prototypes to produce class probabilities (Eq. 2). This process evaluated binary few-shot classification without fine-tuning the meta-parameters *θ*, allowing the assessment of the model’s generalization to unseen studies. For each shot value *K* and each cohort, we repeated such evaluation procedure 100 times to obtain performance statistics.

### 2.5 Baseline models for comparison

To comprehensively access the contribution of the learned STUNT embedding, we constructed several baseline methods that were trained on only the held-out studies, using the same repeated episodic splits as STUNT. This allows us to separate the benefits of the pretrained representations on generalization to unseen studies. Additionally, we built a Logistic Regression and a Random Forest on top of the STUNT embedding to evaluate whether the learned representations is useful for other classifiers as well. These baseline models are described below.

#### Prototypical Network on raw features (Raw Prototype)

This model performs prototype-based classification as STUNT but directly in the raw feature space without leveraging the meta-learned embedding. Therefore, for each evaluation procedure, prototypes are computed as the mean of the support samples in *S* and query samples in *Q* are then assigned to the nearest prototype. This baseline isolates the contribution of the learned embedding by examining the prototype-based classification strategy alone.

#### Few-shot Logistic Regression on STUNT embeddings (Few-shot LR on STUNT)

Instead of the prototype-based classification, this model trains a Logistic Regression classifier was trained on STUNT-derived embeddings of the support set *S* and evaluated on embeddings of the query set *Q*.

#### Few-shot Random Forest on STUNT embeddings (Few-shot RF on STUNT)

Similarly to Few-shot LR on STUNT, a Random Forest classifier was trained on top of the STUNT-derived embeddings of the support set *S* and evaluated on the query set *Q*.

#### Few-shot Logistic Regression (Few-shot LR)

A Logistic Regression with L2 regularization (C=1.0) trained from scratch using only the support set *S* and evaluated using the query set *Q*. This model represents a traditional linear machine learning baseline trained with raw features under the same data constraints as STUNT.

#### Few-shot Random Forest (Few-shot RF)

A Random Forest classifier with 100 trees and maximum depth of 3, trained from scratch similarly as Few-shot LR using only the support set *S* and evaluated using the query set *Q*. This model represents a traditional non-linear machine learning baseline trained with raw features under the same data constraints as STUNT.

#### Full-data Logistic Regression (Full-data LR)

To establish an upper bound on achievable performance of Logistic Regression, we trained another LR classifier with L2 regularization (C=1.0) on all available labeled data from the study except the query set, which combines the samples from the support set *S* and remaining set *R*.

#### Full-data Random Forest (Full-data RF)

Likewise, to establish an upper bound of RF when abundant data is available, we trained another RF classifier with 100 trees and maximum depth of 5 on all available labeled data from the study except the query set.

### 2.6 Evaluation metrics and statistical tests

Model performance was evaluated using area under the ROC curve (AUC-ROC), macro F1, and balanced accuracy. These metrics provide complementary assessments of classification performance, particularly under potential class imbalance.

**Area Under the ROC Curve (AUC-ROC)** quantifies a classifier’s ability to discriminate between classes across all decision thresholds. An AUC of 0.5 indicates random performance, whereas an AUC of 1.0 indicates perfect discrimination. Formally, the AUC corresponds to the area under the ROC curve, which plots the True Positive Rate (TPR) against the False Positive Rate (FPR) as the classification threshold varies.

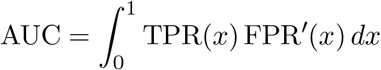

**Macro F1** calculates the unweighted mean of the F1 scores for each class, giving equal importance to all classes regardless of their sample size. For binary classification, this reduces to:

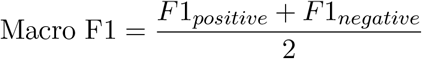

For each class *c*, the F1 score is the harmonic mean of its precision and recall.

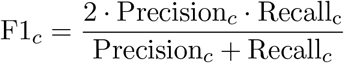

where precision and recall for class *c* are given by:

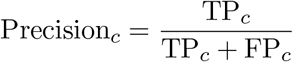

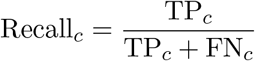

Here TP*_c_*, FP*_c_*, and FN*_c_* denote the numbers of true positives, false positives, and false negatives computed with respect to class *c*.

**Balanced Accuracy** measures classification performance by averaging the recall obtained for each class, thereby accounting for potential class imbalance. For binary classification, balanced accuracy is defined as the average of sensitivity and specificity.

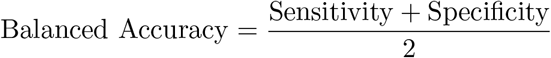

To assess the significance of the observed performance differences between STUNT-based models and other baseline models, we employed the Wilcoxon signed-rank test to compare comparing each STUNT-based method with its raw-feature counterpart, with a two-sided alternative hypothesis testing whether one method systematically outperforms the other across evaluation iterations. Statistical significance was assessed at *α* = 0.05.

## 3 Results

### 3.1 Overview of the five metagenomic studies used for model evaluation

We evaluated model performance using five publicly available gut microbiome datasets, which are all whole-genome shotgun metagenomic datasets selected based on the quality control, metadata and species-level abundance data provided from GMrepo v2 database. As shown in Figure 2A, the associated diseases of the chosen studies included rheumatoid arthritis (RA; PRJNA356102, n=168), gestational diabetes mellitus during pregnancy (GDM, Pregnant; PRJNA401977, n=144), non-alcoholic fatty liver disease (NAFLD; PRJNA373901, n=86), diabetes mellitus, type 1 (T1D; PRJNA289586, n=53), and inflammatory bowel diseases (IBD; PRJEB7949, n=40). The sample sizes ranged from 40 (IBD cohort) to 168 (RA cohort) and class distributions were moderately balanced across studies. Samples were represented by CLR- and Z-score-transformed relative abundances of 177 species-level taxa, selected according to the low-abundance filtering result from the meta-training data.

**Figure 2:**
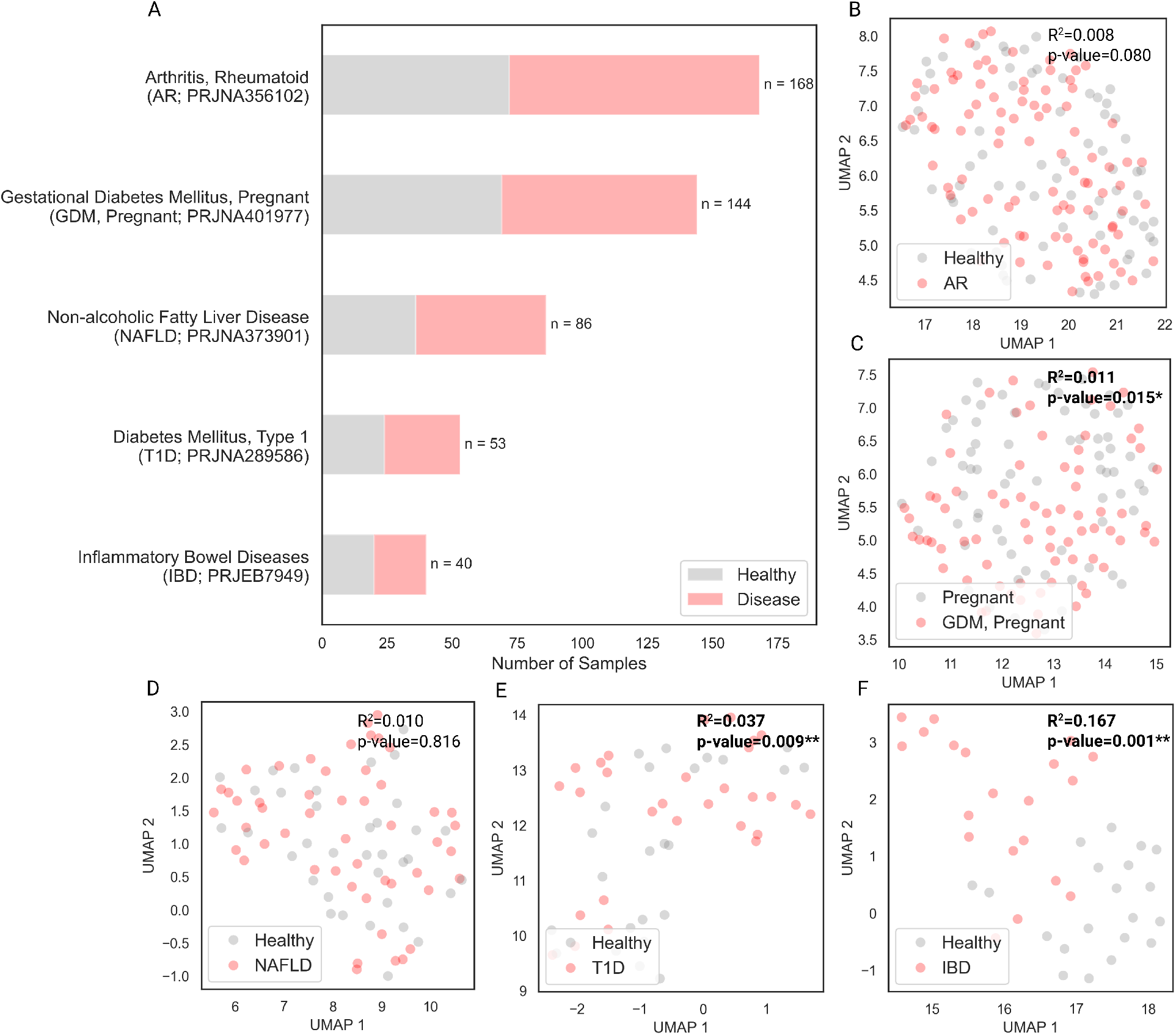
Overview of the five held-out metagenomic cohorts used for model evaluation.. (A) Summary of the selected cohorts, including project ID, associated disease, and total sample size, stratified by the health status. (B-F) Two-dimensional UMAP projections and PERMANOVA results of the pre-processed species-level sample profiles for each study. For each study, the R² value represents the proportion of total variance explained by the phenotype factor according to PERMANOVA and the associated p-values were derived from 999 permutations with significance levels: * *p <* 0.05, ** *p <* 0.01, *** *p <* 0.001.

Dimension reduction using UMAP and PERMANOVA revealed varying degrees of class separation between the disease and control samples across the five cohorts (Figure 2 B-F). The PERMANOVA revealed that among the three studies with statistically significant separation, IBD cohort exhibited the strongest effect (R² = 0.167, p-value = 0.001; Figure 2F), followed by T1D (R² =0.037, p-value = 0.009; Figure 2E) and GDM (R² = 0.011, p-value = 0.015; Figure 2C). In contrast, RA (R² = 0.008, p-value = 0.080; Figure 2B) and NAFLD cohort (R² = 0.010, p-value = 0.816; Figure 2D) exhibited no significant class separation.

### 3.2 STUNT offers no consistent advantages for few-shot microbiome-based disease classification

To assess whether the STUNT embedding improve microbiome-based disease classification under limited sample sizes, we evaluated the classification performance of STUNT and other baseline models using the sample microbial profiles and disease labels of the five chosen studies. Together, the compared models included six few-shot methods: STUNT, Raw Prototype, Few-shot LR on STUNT, Few-shot RF on STUNT, Few-shot LR, and Few-shot RF. Additionally, we included two full-data models, Full-data LR and Full-data RF, which were trained on all available non-query samples for upper bound estimation.

To examine the models’ overall performance across the five held-out studies with varying numbers of available support samples, we constructed the learning curves to compare all methods across the tested shot numbers from *K*=1 to *K*=10. As shown in Figure 3, at *K*=1 STUNT achieved the best performance among few-shot methods, with a mean AUC-ROC of 0.605. Few-shot RF and LR applied on top of the STUNT-derived embeddings followed closely, with mean AUC-ROC of 0.593 and 0.588, respectively. In contrast, methods trained directly on raw features performed slightly worse overall: few-shot LR led this group with an AUC–ROC of 0.586, followed by Raw Prototype at 0.580 and few-shot RF at 0.574. This initial advantage of STUNT-based methods suggests that the learned embedding provides a small benefit when only a single support example per class is available.

**Figure 3:**
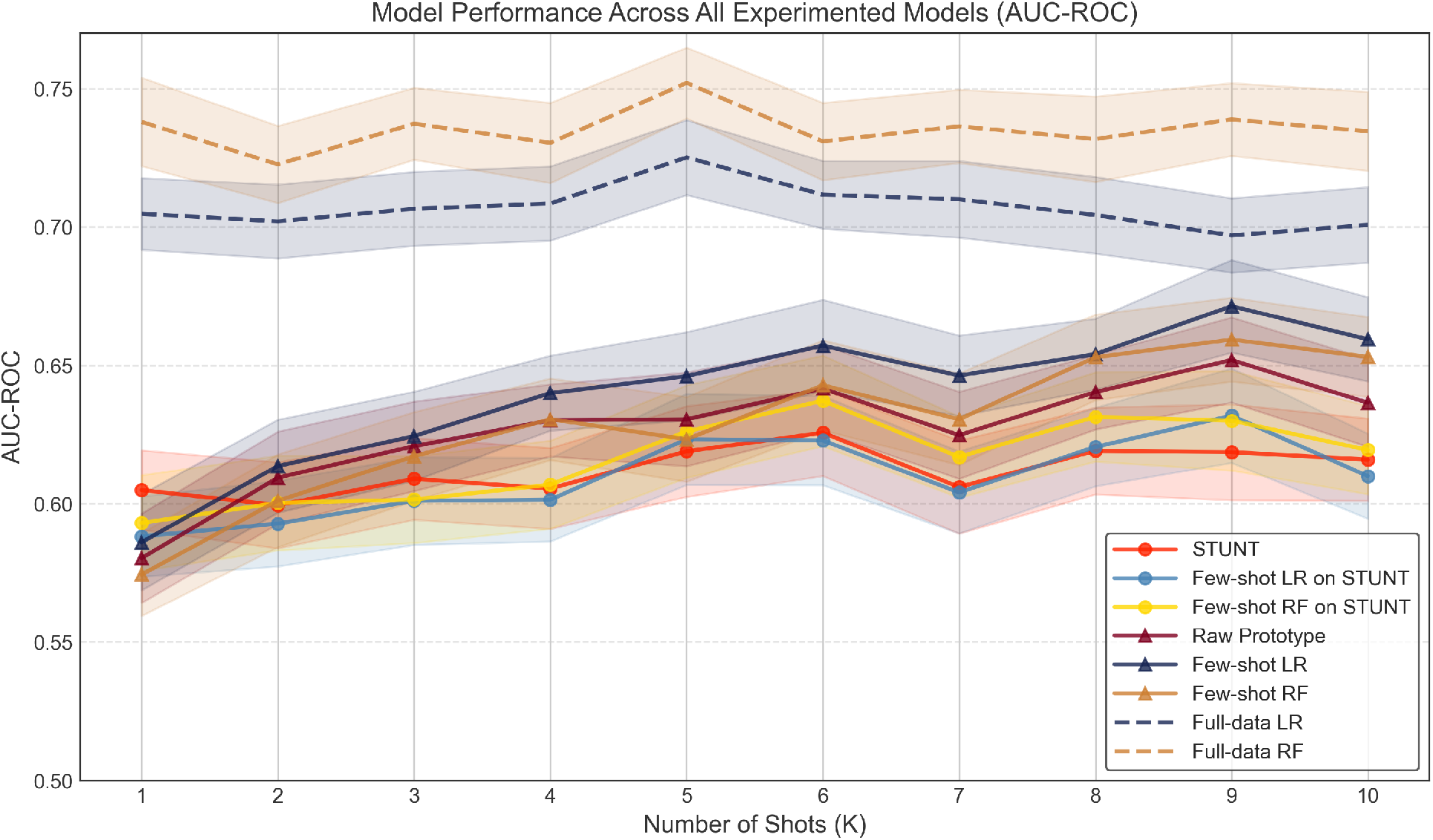
Classification AUC-ROC of STUNT and baseline models using varying shot sizes (K = 1 to 10) across five held-out cohorts. All methods trained under the same few-shot scenario (only on the support set *S* with 2*K* samples) are represented by solid lines, which include six models: STUNT, Few-shot LR on STUNT (Few-shot Logistic Regression on STUNT-derived embeddings), Few-shot RF on STUNT (Few-shot Random Forest on STUNT-derived embeddings), Raw Prototype (Prototypical network on raw features), Few-shot LR (Few-shot Logistic Regression), and Few-shot RF (Few-shot Random Forest). Among these six models, models utilizing STUNT-derived embeddings are indicated by circular markers, while those operating on raw features are indicated by triangular markers. Dashed lines indicate the two baselines trained on all available non-query samples: Full-data LR (Full-data Logistic Regression) and Full-data RF (Full-data Random Forest), serving as upper-bound references. For each k, the AUC-ROC values were averaged across the five studies within each iteration. The means are shown with the dots and the shaded regions represent the 95% confidence intervals across 100 evaluation iterations for each model.

However, this advantage rapidly diminished and reversed with increasing support set size. For instance, by *K*=5, Few-shot LR using raw features had become the best-performing few-shot model, achieving an AUC–ROC of 0.646, followed by Raw Prototype at 0.631, while STUNT dropped to the lowest position among the few-shot methods at 0.619. At K=10, this performance gap was even more pronounced as the model using only raw features continued to improve, reaching 0.659 for Few-shot LR, 0.653 for Few-shot RF, and 0.636 for Raw Prototype. In contrast, all STUNT-based approaches plateaued, with performance remaining around 0.61–0.62. This performance shift is also clearly captured by the ranking trajectory shown in Supplementary Figure S1, which tracked the mean performance ranking for all few-shot models across *K*=1 to 10.

As anticipated, Full-data RF and Full-data LR consistently outperformed all few-shot methods across all tested *K* values. Averaged across all *K* values and studies, Full-data RF achieved a higher overall AUC-ROC of 0.74 than Full-data LR, which achieved 0.71, whereas the best-performing few-shot method, Few-shot LR, reached 0.64. Results for F1 score and balanced accuracy followed similar patterns, with STUNT-based methods showing modest advantages only at *K*=1 that diminished or reversed at higher shot sizes (Supplementary Figures S2).

To isolate the contribution of the STUNT-derived embeddings from the choice of classifier across different number of shots (*K*), we performed direct comparisons between equivalent method pairs that differed only in their assess to STUNT embedding function: STUNT versus Raw Prototype, Few-shot LR on STUNT versus Few-shot LR, and Few-shot RF on STUNT versus Few-shot RF (Figure 4). As shown, STUNT improved the overall classification AUC-ROC across the evaluation iterations at *K*=1 for both Prototypical Network and Random Forest approaches (Figure 4A). Specifically, under this extreme condition, while the absolute improvements are modest but statistically significant: STUNT outperformed Raw Prototype by *△*AUC-ROC=+0.024 with a win rate of 66% (*p <* 0.001), and Few-shot RF on STUNT outperformed Few-shot RF by *△*AUC-ROC=+0.019 with a win rate of 60% (*p <* 0.05), indicated in the Figure 4B. However, this advantage turned into disadvantage at *K*=4 for both methods. Using raw features became consistently better for Prototypical Network at *K ≥*6 and at *K ≥*8 for Random Forest. On the other hand, Logistic Regression never gained a statistically significant advantage from STUNT-derived embeddings and using raw features was significantly better for *K ≥*2. These findings suggest that while STUNT meta-learned embedding might provide modest benefit under extreme data scarcity (*K*=1), they impose an information bottleneck that becomes increasingly detrimental as support set size grows. Overall, the narrow window and small magnitude of the advantage (*K*=1), as well as the substantial disadvantage at higher *K* values suggest that STUNT embedding does not offer consistent benefit for few-shot microbiome classification tasks.

**Figure 4:**
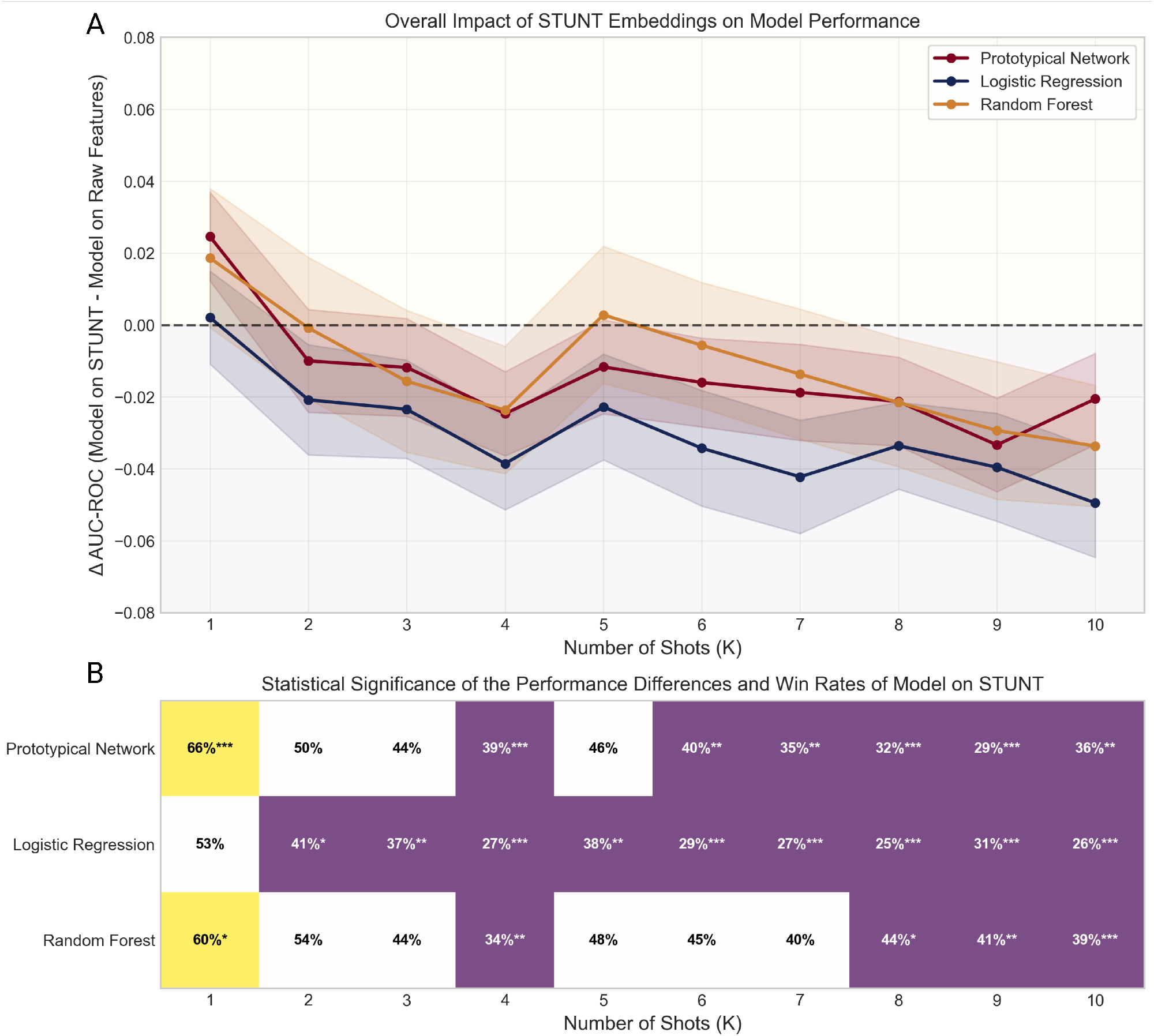
Overall impact of STUNT-derived embeddings on model performance with varying shot sizes (AUC-ROC) (A) Performance difference (*△*AUC-ROC) at each k-shot of the paired classifiers leveraging STUNT-derived embeddings versus raw features: Prototypical Network (STUNT versus Raw Prototype), Logistic Regression (Few-shot LR on STUNT versus Few-shot LR), and Random Forest (Few-shot RF on STUNT versus Few-shot RF). Positive values indicate superior performance achieved by the STUNT-based method over its raw-feature counterpart. Shaded regions indicate 95% confidence intervals computed across 100 iterations. (B) Win rates for STUNT-based methods and statistical significance of the performance differences. The cell values show the proportion of wins by methods which leveraged STUNT-derived embeddings against their raw-feature counterparts across 100 evaluation iterations at each k-shot level. The cell colors show the Wilcoxon rank test results: yellow indicates the STUNT-based method is significantly better, while purple indicates the method only using raw features is significantly better. The significance level are denoted by asterisks: *p¡0.05, **p¡0.01, ***p¡0.001.

### 3.3 Study-wise analysis reveals model performance substantially varies by dataset

To assess whether classification performance varies by dataset, we examined model performance within each of the five studies across different shot sizes (Figure 5). Performance varied substantially across cohorts. Averaging across all *K* values and few-shot methods, IBD achieved the highest AUC-ROC (0.94), followed by T1D (0.64). In contrast, GDM, RA, and NAFLD performed at or below chance (0.48–0.53). Notably, even the full-data methods failed to achieve strong performance in these three cohorts, with Full-data LR performing below chance level (AUC-ROC = 0.32) in NAFLD. F1 score and balanced accuracy showed the same cross-study pattern (Supplementary Figures S3). This performance ranking closely mirrors the PERMANOVA-based class separability of each cohort, with datasets exhibiting larger *R*^2^ values achieving consistently higher classification accuracy.

**Figure 5:**
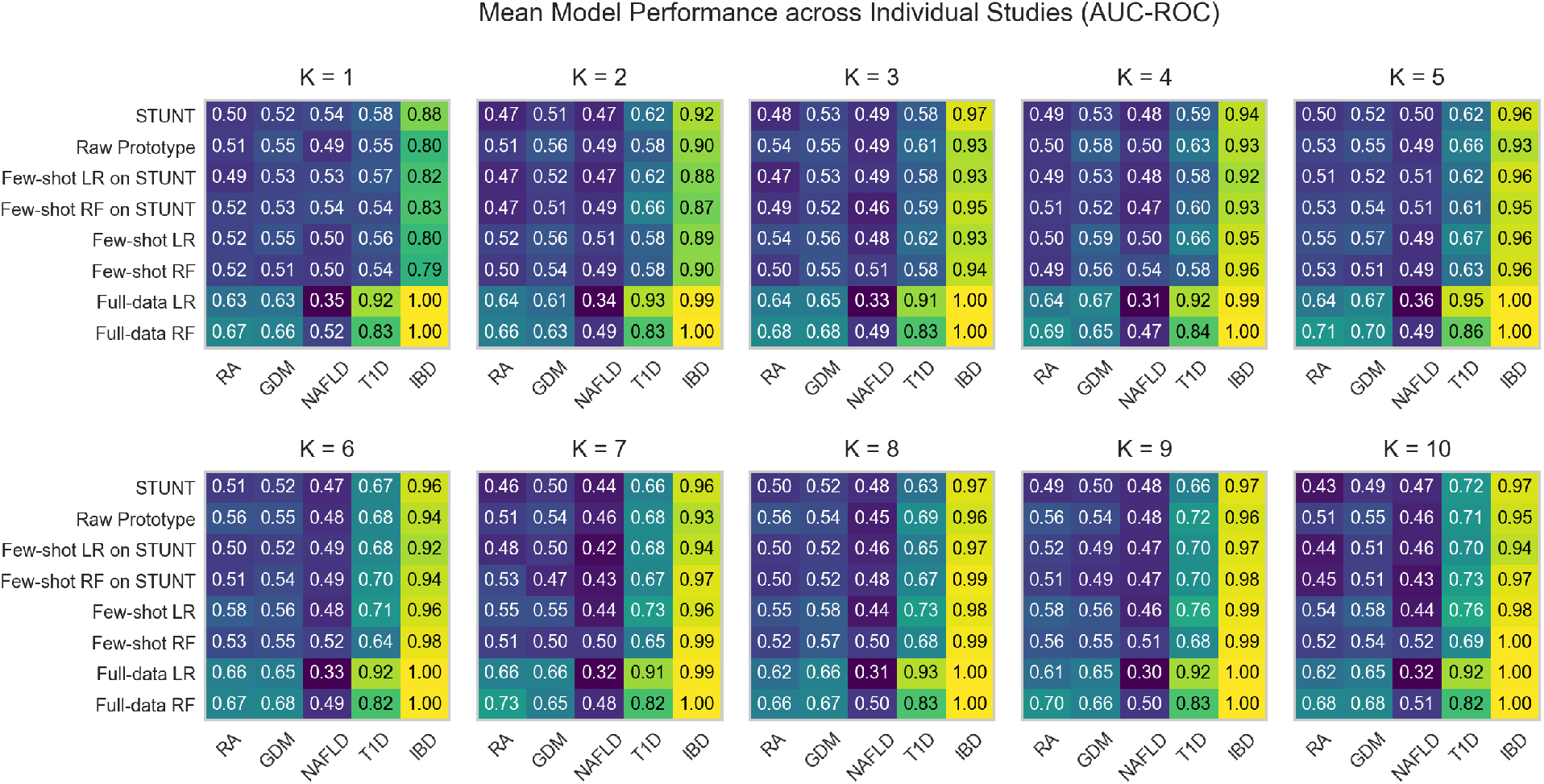
Mean model classification AUC-ROC across individual studies. The heatmaps display the averaged AUC-ROC scores achieved by STUNT and other baseline models (row) within each individual study (column) at each k-shot level across 100 iterations.

Given this cross-study variability, we next assessed whether STUNT-derived embeddings provide study-specific benefits by comparing each STUNT-based method against its raw-feature counterpart within individual studies (Figure 6). For Prototypical Network, STUNT showed consistent and significant improvements only in the IBD cohort, with significant positive differences across 8 out of 10 *K* values. In contrast, the RA, GDM, and T1D cohorts showed the opposite pattern: raw features significantly outperformed STUNT at multiple *K* values, while NAFLD cohort showed no significant difference in either direction. For Logistic Regression and Random Forest trained on STUNT-derived embeddings, neither showed consistent improvement over their raw-feature counterparts across any study. In the IBD cohort, despite STUNT’s advantage for Prototypical Network, both LR and RF on STUNT performed consistently and significantly worse than their raw-feature counterparts at higher shot sizes. Additionally, in the IBD cohort, STUNT showed comparable or worse classification F1 and balanced accuracy than Raw Prototype for *K ≥* 2 (Supplementary Figures S4 and S5). This suggests that STUNT-derived embeddings may improve probability calibration in the IBD cohort but not the actual classification decisions.

**Figure 6:**
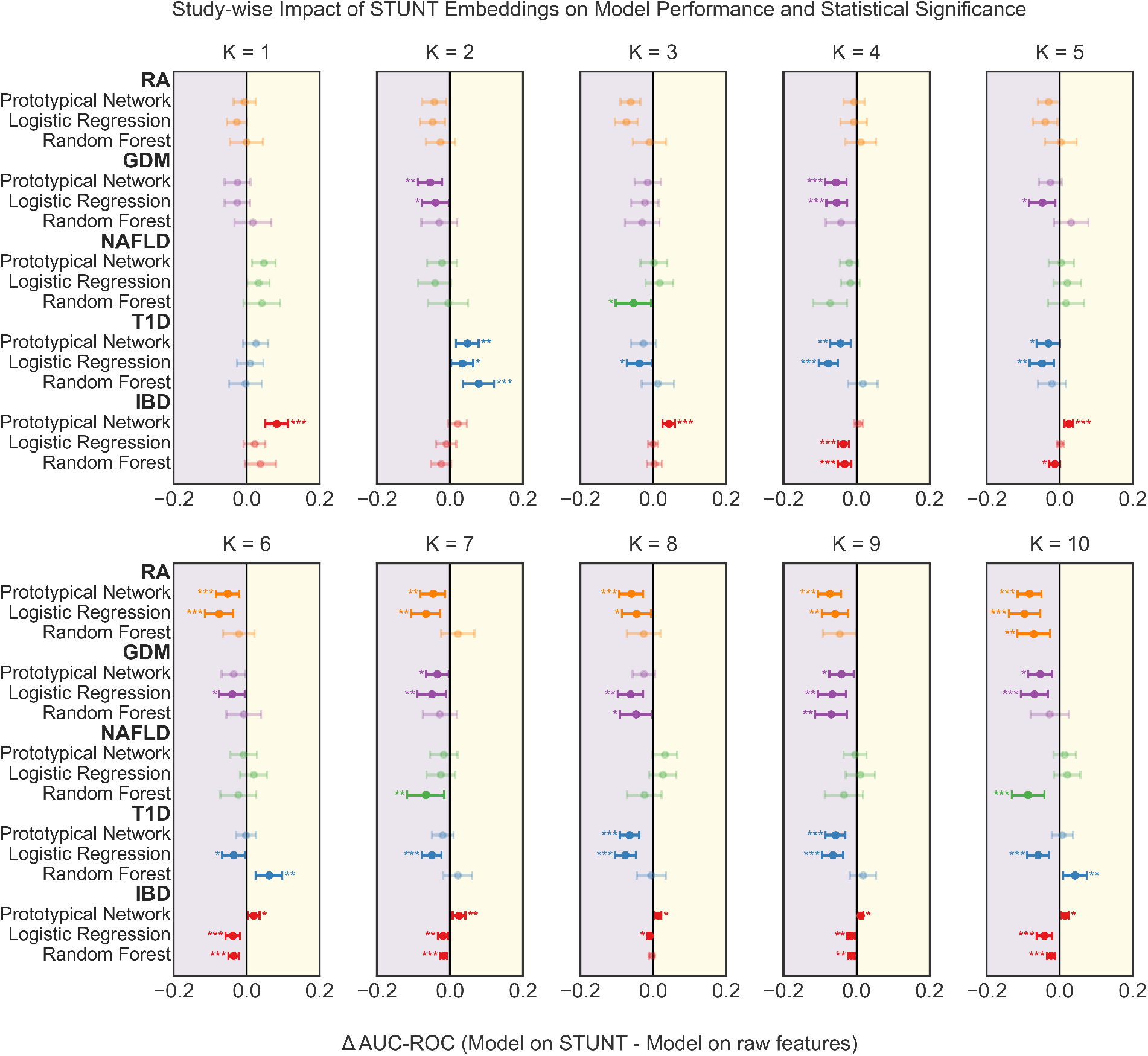
Study-wise impact of STUNT derived embeddings on model performance with varying shot sizes (AUC-ROC). Each panel shows the mean performance difference (*△*AUC-ROC) between STUNT-based methods and their raw-feature counterparts for three classifier types: Prototypical Network (STUNT versus Raw Prototype), Logistic Regression (Few-shot LR on STUNT versus Few-shot LR), and Random Forest (Few-shot RF on STUNT versus Few-shot RF) within each individual study at each k-shot level across 100 iterations, with horizontal error bars showing 95% confidence intervals. Positive values indicate the STUNT-based model outperforms the raw-feature model and vice versa. The significance levels from Wilcoxon signed-rank tests are denoted by asterisks: *p¡0.05, **p¡0.01, ***p¡0.001, with faded points representing non-significant differences.

## 4 Discussion

In this study, we evaluated whether STUNT (Few-shot Tabular Learning with Self-generated Tasks from Unlabeled Tables), a framework that integrates self-supervised representation learning with metric-based meta-learning, can exploit shared microbial structure across publicly available metagenomic cohorts to improve microbiome-based disease classification under few-shot settings. Specifically, we compiled and preprocessed more than 5,000 metagenomic samples across 57 curated cohorts from GMrepo v2, one of the largest human microbiome study databases. During meta-training, we used the species-level metagenomic sample profiles from 52 cohorts to optimize STUNT embedding using self-generated tasks, with various sizes of support set (shot number *K* ranging from 1 to 10). During meta-testing, we systematically compared the performance of Prototypical Network, Logistic Regression, and Random Forest with and without leveraging STUNT-derived embeddings across five previously unseen disease cohorts. Contrary to expectations based on the success of meta-learning in other domains, we found no consistent performance advantage from using the STUNT-derived embeddings. Though STUNT did provide a small measurable benefit when *K* = 1, this advantage rapidly disappeared and reversed with additional labeled samples.

Since STUNT integrates both self-supervised representation learning and meta-learning components, a key open question is which component further improvements should focus on. Our results, together with findings from the current literature, suggest that the primary limitation lies more in the self-supervised representation learning part rather than in the meta-learning framework. In particular, self-supervised representation learning methods are designed to capture broad ecological patterns from heterogeneous unlabeled microbiome data, which may inadvertently discard fine-grained, cohort- or disease-specific signals that are critical for downstream disease prediction.

In our study, this is evidenced by the rapid performance reversal observed between STUNT and Raw Prototype models. STUNT outperforms Raw Prototype in the extremely low-data regime (K=1), indicating that the meta-learned embedding provides meaningful benefits for task initialization and fast adaptation under extreme data scarcity. However, as more labeled samples become available, Raw Prototype increasingly outperforms STUNT, suggesting that the embedding provided diminishing returns as task-specific information can be more effectively learned directly from labeled data.

Broadly, this interpretation is also consistent with recent literature showing that the benefits of leveraging broad self-supervised learning for microbiome-based classification are highly context-depend. In particular, the magnitude of such benefits seems to be inversely associated with the downstream task complexity. For example, substantial improvements have been reported for relatively low-complexity tasks, such as biome classification [13], and moderate gains for intermediate tasks, such as age and BMI prediction [11]. In contrast, similar to our findings, only incremental improvements have been reported for tasks like microbiome-based disease classification [11, 12]. Such high-complexity tasks likely involve much weaker, more heterogeneous and context-specific microbial signals, making them less amenable to broadly learned representations. For such settings, targeted supervised pretraining within the same disease context has demonstrated clearer improvements [14], suggesting that representations more specifically aligned with the downstream domain may be necessary to achieve substantial performance gains.

Interestingly, Prototypical Network and Random Forest both benefited modestly from STUNT-derived embeddings at *K* = 1, with their raw-feature counterparts only becoming significantly superior with *K ≥* 4. In contrast, Logistic Regression appeared to gain no advantage from the meta-learned embeddings under any condition. This discrepancy likely reflects the mismatch between the structure of the meta-learned embedding space and the inductive biases and capacity of Logistic Regression. The STUNT embedding space was optimized specifically for prototype-based non-linear structure to support Prototypical Network, making it still potentially exploitable by Random Forest but not compatible with the linear decision boundary of Logistic Regression. Additionally, using the default setting, STUNT produce 1024-dimensional embeddings, which substantially increases the number of parameters Logistic Regression must estimate, making it more prone to overfitting in few-shot settings. In comparison, Prototypical Network avoids this issue by relying on distance-based classification rather than parameter estimation, while Random Forest can also mitigates it through localized feature splits. Although reducing the default embedding dimensionality could alleviate overfitting, it would further increase the risk of discarding already subtle microbiome signals that are difficult for self-supervised frameworks to capture.

Beyond the limitations of leveraging self-supervised representation learning for microbiome-based disease classification, we identified a more fundamental constraint: for certain cohorts, gut microbiome taxonomic profiles may not serve as strong indicators for the disease status. In our held-out cohorts, PERMANOVA revealed statistically significant separation only for the IBD (*R*^2^ = 0.167), T1D (*R*^2^ = 0.037), and GDM (*R*^2^ = 0.011) cohorts, while RA (*R*^2^ = 0.008) and NAFLD (*R*^2^ = 0.010) cohorts exhibited no significant compositional differences between the case and control samples. The effect sizes also appear to be indicative for the later classification performance. Specifically, while few-shot methods achieved high classification performance for the IBD and modest performance for the T1D cohort, they struggled with the GDM, RA, and NAFLD cohorts. With the access to all available samples, the two full-data classifiers improved the classification of GDM and RA, but still failed with below-chance performance in the NAFLD cohort. These findings highlight the problem of low signal-to-noise ratio in disease classification tasks with microbiome profiles. Also, this limitation seems to impose a hard upper bound on the achievable accuracy, rather than the size of available data and the choice of learning methods. These observations confirm that disease-associated microbial alterations are often subtle, heterogeneous, and easily overshadowed by host, environmental, and technical variability.

Taken together, our findings outline several priorities for future work on developing pretrained models for microbiome-based disease classification. First, improved prediction performance will likely require representation learning strategies that preserve cohort- and disease-relevant variation rather than relying solely on broadly self-supervised objectives. Approaches such as disease-specific pretraining, cohort-stratified learning, and integration with other host metadata may be helpful for capturing the subtle signals in the gut microbiome taxonomic profiles. Second, our results underscore the importance of considering the intrinsic biological signal strength when evaluating methodological advances in microbiome machine learning. Recognizing these limits will be essential for developing more realistic expectations and more targeted modeling strategies for microbiome-based disease prediction.

## 5 Data Availability Statement

The parsed and processed metagenomic datasets, along with the code used in this work are available at https://github.com/Chengyao-Peng/microbiome_meta_learning.

## Notes

### Competing Interest Statement

The authors have declared no competing interest.

### Summary of Updates

This version adds the code availability section and corrects a LaTeX bibliography formatting error

