## Supplementary materials for "Evaluating Few-Shot Meta-Learning using STUNT for Microbiome-Based Disease Classification"

Table S1: **Classifier hyperparameters and their search ranges.** Both classifiers were tuned via Bayesian optimization (Optuna v4.4.0) with 30 trials per inner fold.

| Classifier | Hyperparameter | Range | Sampling |
|---|---|---|---|
| LDA | Solver | svd, lsqr, eigen | Categorical |
| | Shrinkage[a] | None, auto, [0.0, 1.0] | Categorical / Uniform |
| RF | Number of trees | 50–300 | Uniform |
| | Maximum depth | 3–20 | Log-uniform |
| | Minimum samples to split | 2–10 | Uniform |
| | Minimum samples per leaf | 1–4 | Uniform |

[a] Only applicable when solver $\in$ {lsqr, eigen}. “auto” uses the Ledoit–Wolf estimator.

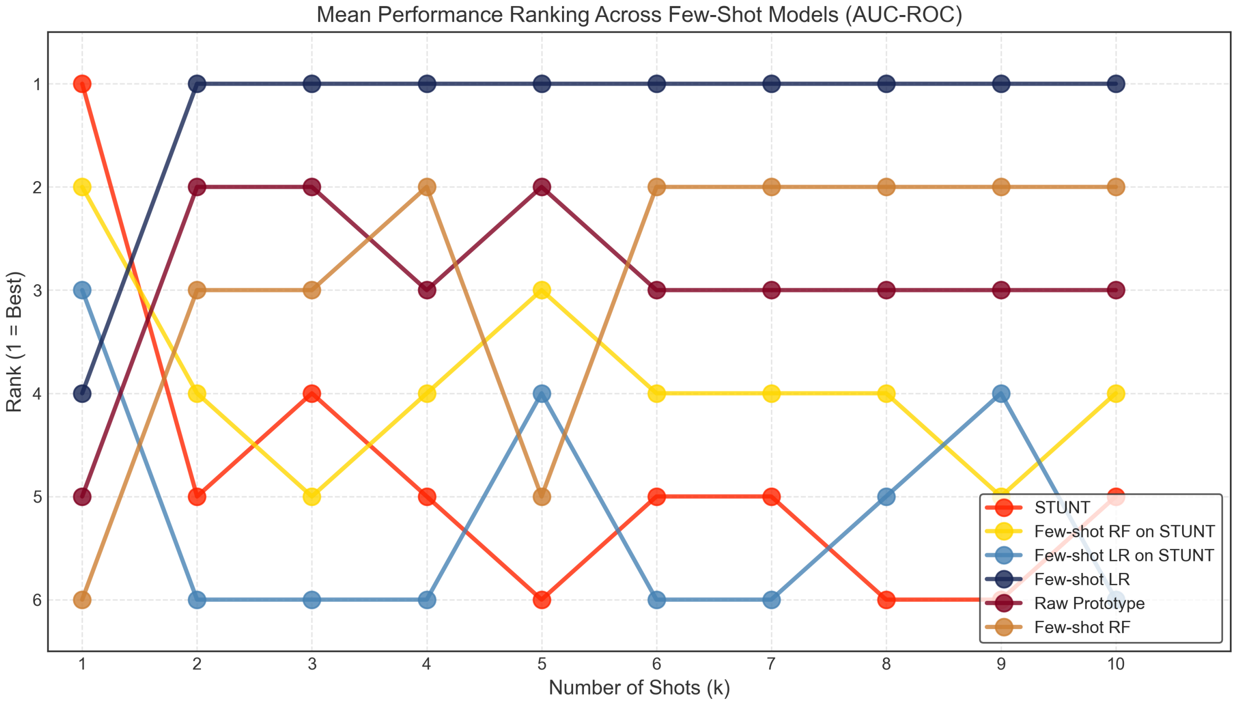


Figure S1: **Mean performance ranking trajectories of few-shot models (AUC-ROC) using varying numbers of support samples (k)**. The rankings were determined by mean AUC-ROC averaged across all five studies at each k, with rank 1 indicating the best-performing method and 6 the worst-performing method.

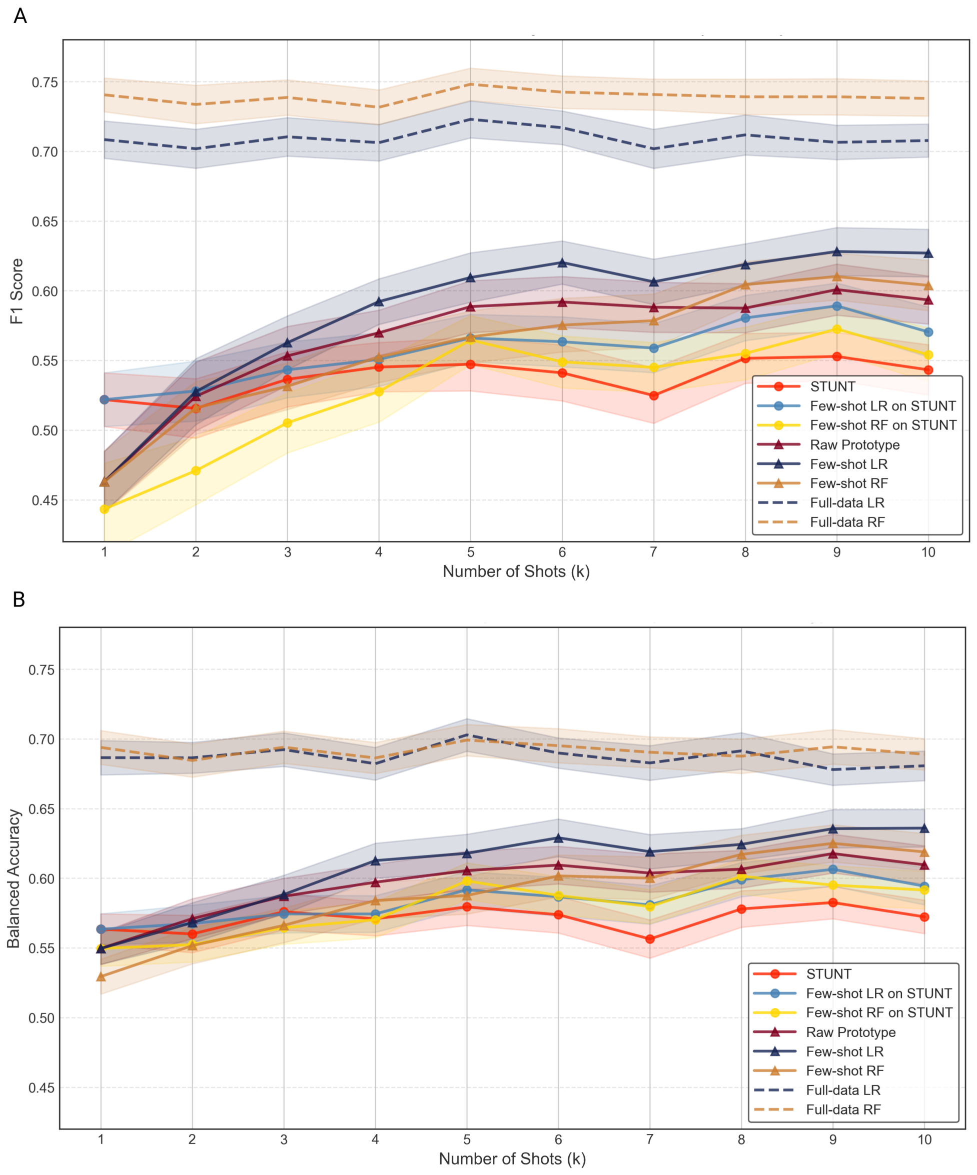


Figure S2: **Classification macro F1 and balanced accuracy of STUNT and baseline models with varying shot sizes ($K$ = 1 to 10) across five held-out cohorts.** Panel A shows macro F1 scores and Panel B shows balanced accuracy. All methods trained under the same few-shot scenario (only on the support set $S$ with 2k samples) are represented by solid lines, which include six models: STUNT, Few-shot LR on STUNT (Few-shot Logistic Regression on STUNT-derived embeddings), Few-shot RF on STUNT (Few-shot Random Forest on STUNT-derived embeddings), Raw Prototype (Prototypical network on raw features), Few-shot LR (Few-shot Logistic Regression), and Few-shot RF (Few-shot Random Forest). Among these six models, Models utilizing STUNT meta-learned embeddings are indicated by circular markers, while those operating on raw features are indicated by triangular markers. Dashed lines indicate the two baselines trained on all available non-query samples: Full-data LR (Full-data Logistic Regression) and Full-data RF (Full-data Random Forest), serving as upper-bound references. For each k, both metrics were averaged across the five studies within each iteration. The means are shown with the dots and the shaded regions represent the 95% confidence intervals across 100 evaluation iterations for each model.

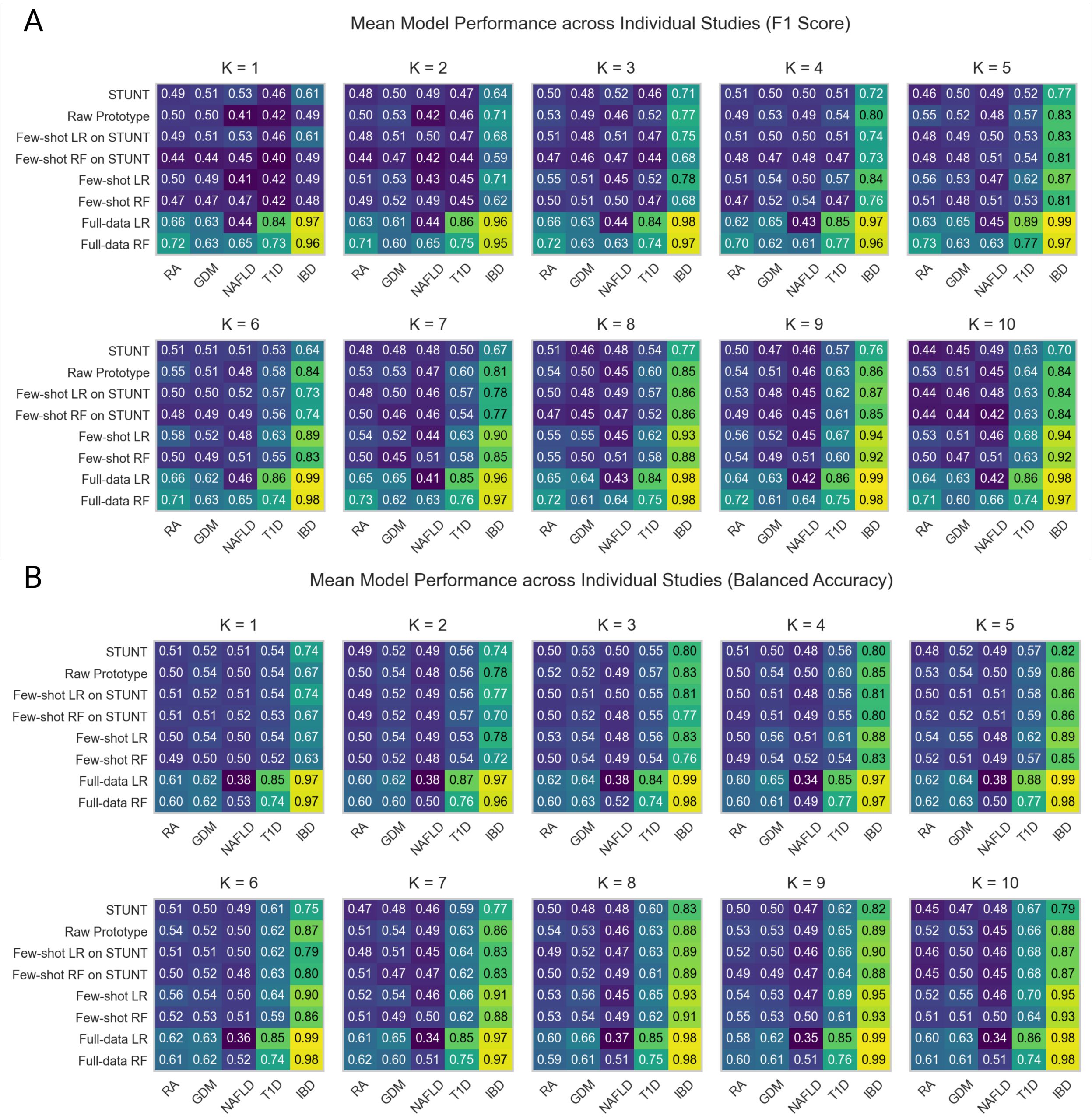


Figure S3: **Mean model F1 score and balanced accuracy across individual studies.** The heatmaps display the averaged F1 scores in Panel A and balanced accuracy in Panel B, achieved by STUNT and other baseline models (row) within each individual study (column) at each k-shot level across 100 iterations.

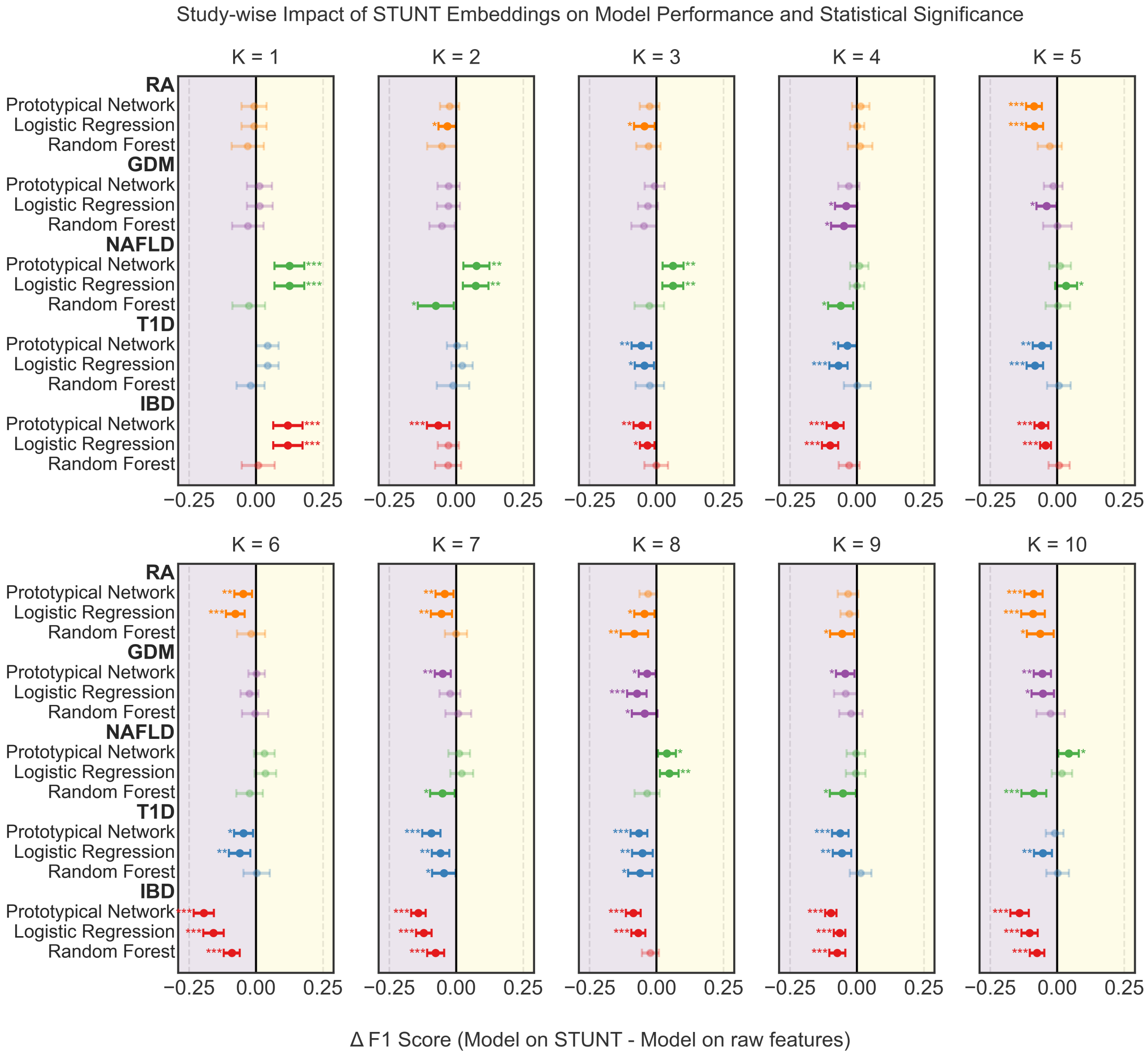


Figure S4: **Study-wise impact of STUNT embeddings on model performance with varying shot sizes (F1 Score).** Each panel shows the mean performance difference (△F1 Score) between STUNT-based methods and their raw-feature counterparts for three classifier types: Prototypical Network (STUNT versus Raw Prototype), Logistic Regression (Few-shot LR on STUNT vs Few-shot LR), and Random Forest (Few-shot RF on STUNT versus Few-shot RF) within each individual study at each k-shot level across 100 iterations, with horizontal error bars showing 95% confidence intervals. Positive values indicate the STUNT-based model outperforms the raw-feature model and vice versa. The significance levels from Wilcoxon signed-rank tests are denoted by asterisks: *p¡0.05, **p¡0.01, ***p¡0.001, with faded points representing non-significant differences.

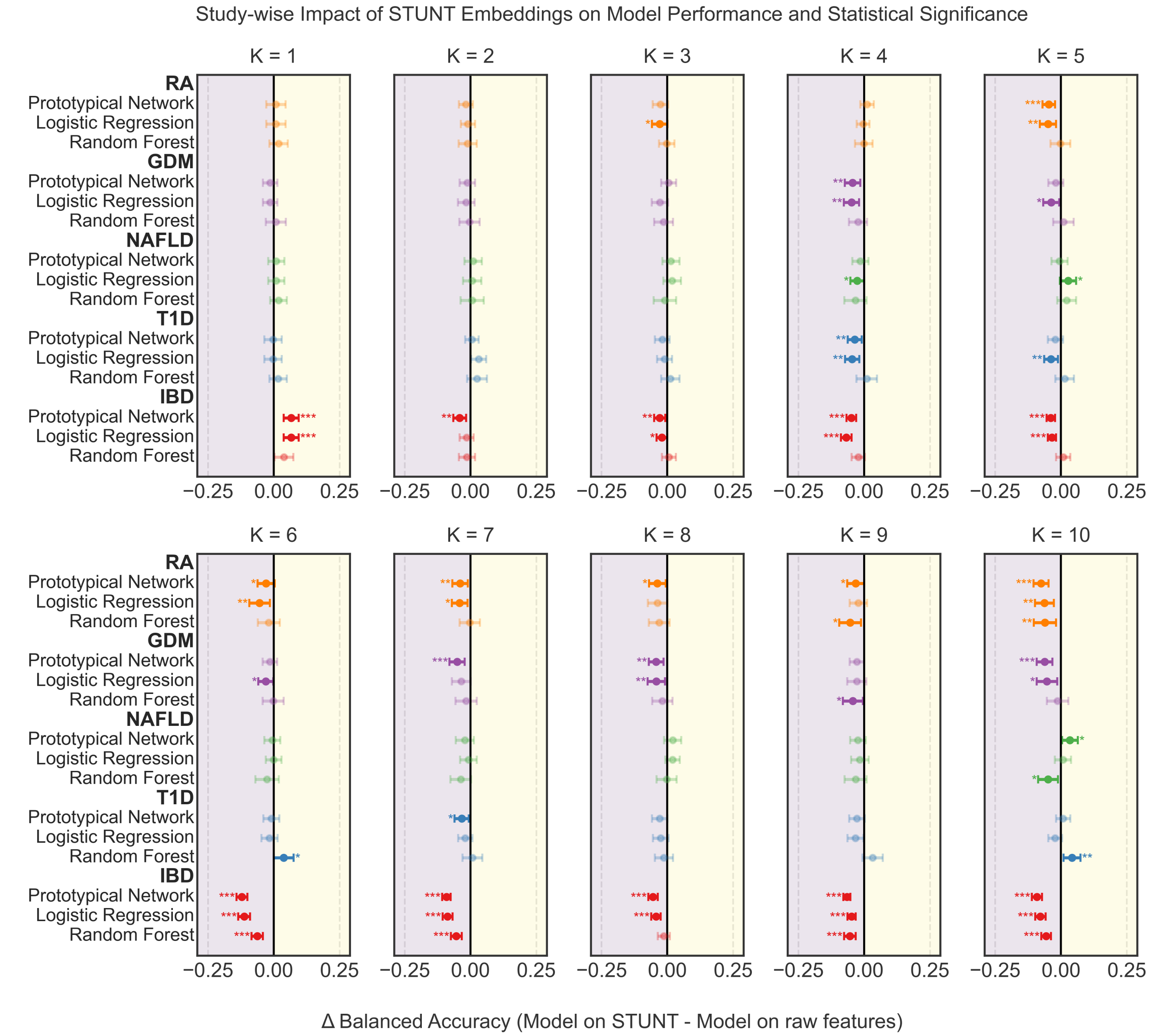


Figure S5: **Study-wise impact of STUNT-derived embeddings on model performance with varying shot sizes (balanced accuracy).** Each panel shows the mean performance difference (△balanced accuracy) between STUNT-based methods and their raw-feature counterparts for three classifier types: Prototypical Network (STUNT versus Raw Prototype), Logistic Regression (Few-shot LR on STUNT versus Few-shot LR), and Random Forest (Few-shot RF on STUNT versus Few-shot RF) within each individual study at each k-shot level across 100 iterations, with horizontal error bars showing 95% confidence intervals. Positive values indicate the STUNT-based model outperforms the raw-feature model and vice versa. The significance levels from Wilcoxon signed-rank tests are denoted by asterisks: *p¡0.05, **p¡0.01, ***p¡0.001, with faded points representing non-significant differences.
